# Transcriptional Profiling of Somatostatin Interneurons in the Spinal Dorsal Horn

**DOI:** 10.1101/215657

**Authors:** Alexander Chamessian, Michael Young, Yawar Qadri, Temugin Berta, Ru-Rong Ji, Thomas Van De Ven

**Affiliations:** Department of Anesthesiology, Duke University Medical Center, Durham, North Carolina, 27710, USA; Department of Neurobiology, Duke University Medical Center, Durham, North Carolina, 27710, USA; Medical Scientist Training Program, Duke University School of Medicine, North Carolina, 27710, USA; Department of Pharmacology and Cancer Biology, Duke University Medical Center, Durham, North Carolina, 27710, USA; Pain Research Center, Department of Anesthesiology, University of Cincinnati Medical Center, Cincinnati, Ohio, 45267, USA

## Abstract

The spinal dorsal horn (SDH) is comprised of distinct neuronal populations that process different somatosensory modalities. Somatostatin (SST)-expressing interneurons in the SDH have been implicated specifically in mediating mechanical pain. Identifying the transcriptomic profile of SST neurons could elucidate the unique genetic features of this population and enable selective analgesic targeting. To that end, we combined the Isolation of Nuclei Tagged in Specific Cell Types (INTACT) method and Fluorescence Activated Nuclei Sorting (FANS) to capture tagged SST nuclei in the SDH of adult male mice. Using RNA-sequencing (RNA-seq), we uncovered more than 13,000 genes. Differential gene expression analysis revealed more than 900 genes with at least 2-fold enrichment. In addition to many known dorsal horn genes, we identified and validated several novel transcripts from pharmacologically tractable functional classes: Carbonic Anhydrase 12 *(Car12)*, Phosphodiesterase 11A *(Pde11a)*, Protease-Activated Receptor 3 *(F2rl2)* and G-protein Coupled Receptor 26 *(Gpr26). In situ* hybridization of these novel genes revealed differential expression patterns in the SDH, demonstrating the presence of transcriptionally distinct subpopulations within the SST population. Pathway analysis revealed several enriched signaling pathways including cyclic AMP-mediated signaling, Nitric Oxide Synthase signaling, and voltage-gated calcium channels, highlighting the importance of these pathways to SST neuron function. Overall, our findings provide new insights into the gene repertoire of SST dorsal horn neurons and reveal several candidate targets for pharmacological modulation of this pain-mediating population.

**Significance Statement:** Somatostatin(SST)-expressing interneurons in the spinal dorsal horn (SDH) are required for the perception of mechanical pain. Identifying the distinctive genes expressed by SST neurons could facilitate the development of novel, circuit-targeting analgesics. Thus, we applied cell type-specific RNA-sequencing (RNA-seq) to provide the first transcriptional profile of SST neurons in the SDH. Bioinformatic analysis revealed hundreds of genes enriched in SST neurons, including several previously undescribed genes from druggable classes (*Car12, Pde11a, F2rl2* and *Gpr26*). Taken together, our study unveils a comprehensive transcriptional signature for SST neurons, highlights promising candidate genes for future analgesic development, and establishes a flexible method for transcriptional profiling of any spinal cord cell type.

## Introduction

Mechanical pain is one of the chief symptoms in many pathological pain conditions (Costigan et al., 2009b; Cohen and Mao, 2014). Accordingly, understanding the spinal circuits underlying this component of pain perception has been a central aim of preclinical pain research(Peirs and Seal, 2016; Duan et al., 2017). Recent studies employing genetic tools to manipulate specific spinal circuits have greatly expanded our understanding of this question(Duan et al., 2014; Peirs et al., 2015; Cheng et al., 2017; Francois et al., 2017). In this way, it was discovered that a population of Somatostatin-expressing (SST) excitatory interneurons in the superficial dorsal horn is required for mechanical pain, as demonstrated by the complete absence of mechanical pain when SST neurons were ablated(Duan et al., 2014)}. Importantly, other sensory modalities such as thermosensation and innocuous touch were left undisturbed, indicating that SST interneurons play a specific and restricted role. With the functional role of SST neurons now well-established, it would be advantageous to comprehensively characterize the repertoire of genes expressed by SST neurons, thus providing a basis for their unique properties and highlighting potential targets for selective pharmacological manipulation.

RNA-sequencing (RNA-seq) is a powerful tool to uncover the transcriptome of cells and tissues. To date, gene expression studies of the dorsal horn have used RNA isolated from bulk spinal tissue, which represents a mixture of genes expressed by multiple neuronal and non-neuronal cell types(Costigan et al., 2009a; LaCroix-Fralish et al., 2011). To examine the transcriptional profile of a single neuronal population, isolating RNA solely from those cells is necessary(Shin et al., 2014).

To access the transcriptome of specific cell types, various methods have been developed. Dissociation of intact neurons from neural tissue coupled with fluorescence activated cell sorting (FACS) has been used to profile neurons and glia in the CNS, but this method requires relatively harsh protease treatments at warm temperatures, which induces artifactual gene expression signatures due to processing(Lacar et al., 2016; Adam et al., 2017; van den Brink et al., 2017). Moreover, whole cell dissociation biases the eventual sample toward cells that are easier to dissociate. In the case of the dorsal spinal cord from adult mice, standard proteolytic digestion results in a cell suspension that is disproportionately composed of glial cells, with scant viable neurons (YQ, unpublished data).

To obviate the need to dissociate cells, a cell type-specific method called Isolation of Tagged Nuclei from Specific Cell Types (INTACT) was developed (Mo et al., 2015; Steiner and Henikoff, 2015). In this method, nuclei conditionally express a fusion protein comprised of green fluorescent protein (GFP) and the native nuclear membrane protein SUN1 (Lei et al., 2009). Tagged nuclei can then be captured either by immunoprecipitation or fluorescence activated nuclear sorting (FANS) for downstream RNA or DNA analysis. The INTACT method has many benefits. Nuclei are readily obtained from fresh or frozen tissue at cold temperatures by simple mechanical homogenization, which eliminates the concern of processing artifacts and offers the unique possibility to use post-mortem human samples(Grindberg et al., 2013; Lake et al., 2016). Because intercellular connections are destroyed by mechanical dissociation, biases toward specific cell types due to viability or cytoarchitecture are greatly minimized(Lacar et al., 2016). Moreover, it has been demonstrated that the transcriptional signature of nuclear RNA is highly concordant (~95%) with that of whole cells (Grindberg et al., 2013; Lake et al., 2017).

In this study, we optimized INTACT for use on spinal cord tissue to profile the transcriptome of dorsal horn SST neurons. We determined the expression levels of > 13,000 genes, 901 of which were significantly enriched in the SST population compared to all dorsal horn cells. Using *in situ* hybridization and immunohistochemistry, we validated the expression of several novel and highly enriched genes in SST neurons that would make attractive therapeutic targets. Furthermore, we show the SST population is transcriptionally heterogeneous and contains multiple subpopulations.

## Methods and Materials

### Animals

All the animal procedures were approved by the Institutional Animal Care and Use Committee of Duke University. Animal experiments were conducted in accordance with the NIH Guide for the Care and Use of Laboratory Animals. Mouse strains used included *Sst-ires-Cre* (Jax#013044) (Taniguchi et al., 2011) and *R26-CAG-LSL-Sun1-sfGFP-Myc* (Jax # 021039) (Mo et al., 2015). We refer to this line as *Sun1-GFP*^*(fl/fl)*^. To generate SST^GFP^ mice, homozygous *Sst-ires-Cre* males were bred with homozygous Sun1-GFP^(fl/fl)^ females to create compound heterozygous offspring, which were used in all subsequent experiments. To generate control animals for comparison to SST^GFP^, we bred homozygous *Sun1-GFP*^*((fl//fl)*^ with C57BL6/J mice to obtain offspring with *Sun1-GFP*^*(f/+)*^ genotypes. For immunohistochemistry in Figure 3Af-i, we used spinal cord tissue from SST-Tomato mice, which is the product of a cross between homozygotes of the *Sst-ires-Cre* line and Cre-dependent tdTomato reporter line *Ai9* (Jax# 007909) (Madisen et al., 2010). For RNA-seq and microscopy experiments, male mice 8-12 weeks of age were used.

### Nuclei isolation and Fluorescence Activated Nuclear Sorting (FANS)

A dorsal segment from the L3-L5 region from each SST^GFP^ or *Sun1-GFP*^*((fl+)*^ mouse was dissected using small spring scissors and snap frozen on dry ice for later processing. On the day of experiment, the tissue segment was placed in 1 mL of Nuclear Extraction Buffer (NEB) supplemented with RNase and Protease Inhibitors (20 mM Tris HCl pH 8, 5 mM MgCl2, 25 mM KCl, 250 mM Sucrose, 40 U/mL RNasin Plus (Promega), 1 tablet/10 mL Protease Cocktail Mini EDTA-Free (Roche), 1 uM DTT (Sigma), 0.3% NP-40 (Pierce)). Dounce Homogenization (10 strokes Pestle A, 10 strokes Pestle B) was performed to liberate the nuclei using a 2 ml homogenizer (Sigma). The homogenate was filtered through a 50 μm Partec filter (Sysmex) into a regular (not low-binding) 1.7 ml microcentrifuge tube (Axygen), since low-binding tubes create loose pellets that are easily displaced. The filter was washed with an additional 700 μl NEB and the homogenate was then centrifuged for 10 mins at 4°C (500g) to form a loose pellet. The pellet was resuspended with 500 μl of NEB using a p1000 pipettor with regular-bore tips and 10 aspiration/dispense cycles, and filtered through a 20 μm Partec filter into a 5 ml polypropylene tube. DAPI (4’,6-diamidino-2-phenylindole) was added to the sample to a final concentration of 5 ng/ml. We made several modifications to the INTACT procedure by Mo *et al.* (Mo et al., 2015) to simplify the workflow and make the procedure suitable for small spinal cord samples: (1) All volumes and vessels were scaled down to accommodate the smaller size of a dorsal lumbar spinal segment from mouse compared to cortex; (2) Density gradient separation (e.g. Iodixanol) was removed, as we noticed it caused clumping of nuclei and added no additional benefit; (3) FANS, as opposed to bead immunoprecipitation, was used in order to most specifically and cleanly isolate the relatively low number of GFP+ nuclei in the dorsal segment.

FANS was performed using a BD FACSAria II sorter (BD Biosciences) using a pressure of 35 pounds per square inch (psi) and a 100 μm sort nozzle. Gates were established to capture singlet nuclei with high DAPI staining intensity. For SST^GFP^, 5000 DAPI+/GFP+ events were sorted into 350 μl of RNAaqeous Micro (Life Technologies) lysis buffer. For the total nuclei control sample from *Sun1-GFP*^*((fl/+)*^ mice, 5000 DAPI+/GFP-events were isolated in the same manner. FANS data were analyzed using FlowJo 3.0 (FlowJo LLC). For each mouse line, n=3 samples were isolated. Captured samples were placed on ice and immediately processed for RNA isolation.

### RNA library construction and sequencing

RNA isolation was performed using the RNaqeuous Micro kit (Life Technologies) according to the manufacturer’s instructions. DNase digestion was not performed at this step, since the downstream library preparation included a DNase step. For library preparation, all samples were processed at the same time using the SoLo RNA-Seq kit (NuGen Technologies) according to the manufacturer’s instructions in a PCR-clean laminar flow hood. For the PCR amplification step, 17 cycles were used, as this was the optimal number of cycles determined in a prior qPCR optimization assay according to the manfacturer’s instructions. Library concentration was assessed with the Qubit 2.0 fluorometer and dsDNA HS assay (Life Technologies), checked for quality on the Bioanalyzer (Agilent) and then run on the HiSeq 2500 (Illumina) using a 50 base-pair, single-end read protocol.

### Bioinformatic Analysis

RNA-seq data was processed using the TrimGalore toolkit (https://www.bioinformatics.babraham.ac.uk/projects/trim_galore/) which employs Cutadapt {Martin:2011eu} to trim low quality bases and Illumina sequencing adapters from the 3’ end of the reads. Only reads that were 20nt or longer after trimming were kept for further analysis. Reads were mapped to the GRCm38v68 version(Kersey et al., 2016) of the mouse genome and transcriptome using the STAR RNA-seq alignment tool {Dobin:2013fg}. Reads were kept for subsequent analysis if they mapped to a single genomic location. Gene counts were compiled using the HTSeq tool {Anders:2015gf}. For this analysis, we used the standard method of only counting reads that mapped to known exons. Only genes that had at least 10 reads in any given library were used in subsequent analysis. Normalization and differential expression was carried out using the DESeq2 (Love et al., 2014) Bioconductor (Huber et al., 2015) package with the R statistical programming environment. The false discovery rate was calculated to control for multiple hypothesis testing.

Heatmap generation for function classes was performed using the pheatmap package (R). Only coding genes with log2FC > 1, q-value < 0.01, coefficient of variation (CV) < 65% and normalized counts > 20 were included. Pathway analysis was performed using the Ingenuity Pathway Analysis program (Qiagen), using only upregulated genes with q-value < 0.01 and log2FC > 1. Gene set enrichment analysis (Mootha et al., 2003) was performed to identify differentially regulated pathways and gene ontology terms for each of the comparisons performed. Up-regulated genes matching the inclusion criteria above were analyzed using Ingenuity Pathway Analysis (Qiagen).

Gene lists for functional classes were curated from the IUPHAR database (Southan et al., 2016), the Pain Genes database (LaCroix-Fralish et al., 2007), Neuropeptides Database (http://www.neuropeptides.nl/tabel%20neuropeptides%20linked.htm), and Riken Transcription Factor Database (http://genome.gsc.riken.jp/TFdb/).http://genome.gsc.riken.jp/TFdb/).

### In situ Hybridization (ISH) and Immunohistochemistry (IHC)

Animals were deeply anesthetized with isoflurane and transcardially perfused with 4% paraformaldehyde in phosphate-buffered saline (pH 7.4). After perfusion, lumbar spinal cord segments (L3-L5) were removed and postfixed in the same fixative for 2 hours at 4°C. Then, the tissues were cryopreserved in 30% sucrose/PBS solution for at least 24 hours. For immunohistochemistry (IHC), tissues were blocked in blocking/staining buffer consisting of 1X PBS (Life Technologies), 1% Bovine Serum Albumin (BSA, Cell Signaling Technologies) and 0.4% Triton-X 100 (Bio-rad) for 1 hour at room temperature (RT). After blocking, the sections were then incubated overnight with primary antibodies diluted in blocking buffer at 4°C. After washing 3 times in PBS for 5 minutes at RT, sections were incubated with the appropriate secondary antibody for 1 hour at RT followed by 3 washes in PBS for 5 minutes. Before mounting, some sections were counterstained with DAPI (Life Technologies). Slides were then mounted in Prolong Gold (Life Technologies). Primary antibodies were used at the following dilutions: chicken anti-GFP (1:1000, Abcam, ab13970), goat anti-mCAR12 (1:500, R&D Systems,AF2345). Secondary antibodies were as follows: Cy3-conjugated anti-Goat Cy3 (1:1000, Jackson ImmunoResearch), Alexa 488-conjugated anti-Chicken (1:1000, Life Technologies)

For *in situ* hybridization, lumbar spinal cord segments were dissected and post-fixed for 2 hours at 4°C. The spinal cords were cryo-sectioned to 14 um, thaw-mounted onto Superfrost Plus (Fisher Scientific) slides, allowed to dry for 20 minutes at RT, and then stored at −80C. *In situ* hybridization was performed using the RNAscope system (Advanced Cell Diagnostics). Tissue pretreatment consisted of 30 minutes of Protease IV at RT. Following pretreatment, probe hybridization and detection with the Multiplex Fluorescence Kit v2 were performed according to the manufacturer’s protocol. Probes included Mm-Sst-C1 (#404631), Mm-Sst-C2 (#404631-C2), Mm-Pde11a (#481841), Mm-Car12 (#429991), Mm-F2rl2 (#489591), Mm-Gpr26 (#317381), Mm-Nmur2 (#314111), Mm-Grp (#317861). In some cases, *in situ* hybridization was followed by IHC to visualize GFP because the native GFP signal was destroyed by the RNAscope pretreatment. After the RNAscope detection, the tissue sections were processed for IHC as indicated above to reveal GFP fluorescence. Following ISH or combined ISH/IHC, slides were mounted using Prolong Gold.

Fluorescence was detected using an epifluorescence microscope (Nikon Eclipse NiE). Images were taken at 20x and 40x magnification with Z-stacks. Stacked images were combined using the Maximum Intensity Projection function in the Elements software (Nikon). For quantification, all images were taken using the same acquisition settings. Cell counts were performed on the entire dorsal horn for each marker. Three to five sections were selected from 3-4 animals, and the counts from all sections for each animal were averaged. Each first section was selected randomly, and each subsequent section was at least 140 um apart, in order to avoid double-counting. For co-expression quantification, all images in a set underwent the same image post-processing, with a threshold uniformly set on each color channel. A custom script in Matlab (MathWorks) was used to count co-expression events. Images were prepared in Photoshop CC (Adobe).

### Experimental Design and Statistical Analysis

For RNA-seq, samples were prepared from three (n=3) adult, male mice from the SST^GFP^ line and the *Sun1-GFP*^*((fl//fl)*^. For differential gene expression analysis, the log2 fold change between GFP+ nuclei from SST^GFP^ mice and total DAPI+ nuclei from *Sun1-GFP*^*(fl/fl)*^ was determined using the DESeq2 package in R. Statistical significance was determined using the Wald test statistic. Adjusted p-value, which we refer to the q-value, reflects correction of the p-value for multiple comparisons. We considered a gene to be differentially expressed if the log2FC > 1, q-value <0.01, coefficient of variation between replicate < 65% and that all replicates in the SST^GFP^ set had at least 20 normalized counts. We used a q-value cutoff that is more stringent than the conventional q <0.05 in order minimize false positives. For ISH counting, we present the percentage of colocalization events and the standard error of the mean (SEM).

## Results

### Isolation of RNA Specifically from Somatostatin Interneurons in the Dorsal Horn

In order to specifically tag the nuclei of SST neurons in the dorsal horn, we crossed the *Sst-ires-Cre* mouse line by the Cre-dependent INTACT reporter line, *Sun1-GFP*. We hereafter refer to offspring of this cross as SST^GFP^. In lumbar spinal cord sections from these mice, GFP-labeled nuclei were present in the dorsal horn, with the most abundant distribution in the superficial laminae (I-II) and some sparse expression in the deeper dorsal horn (III-V) (Figure. 1Aa), consistent with the previously reported expression of SST interneurons in this region.

**Figure 1.**
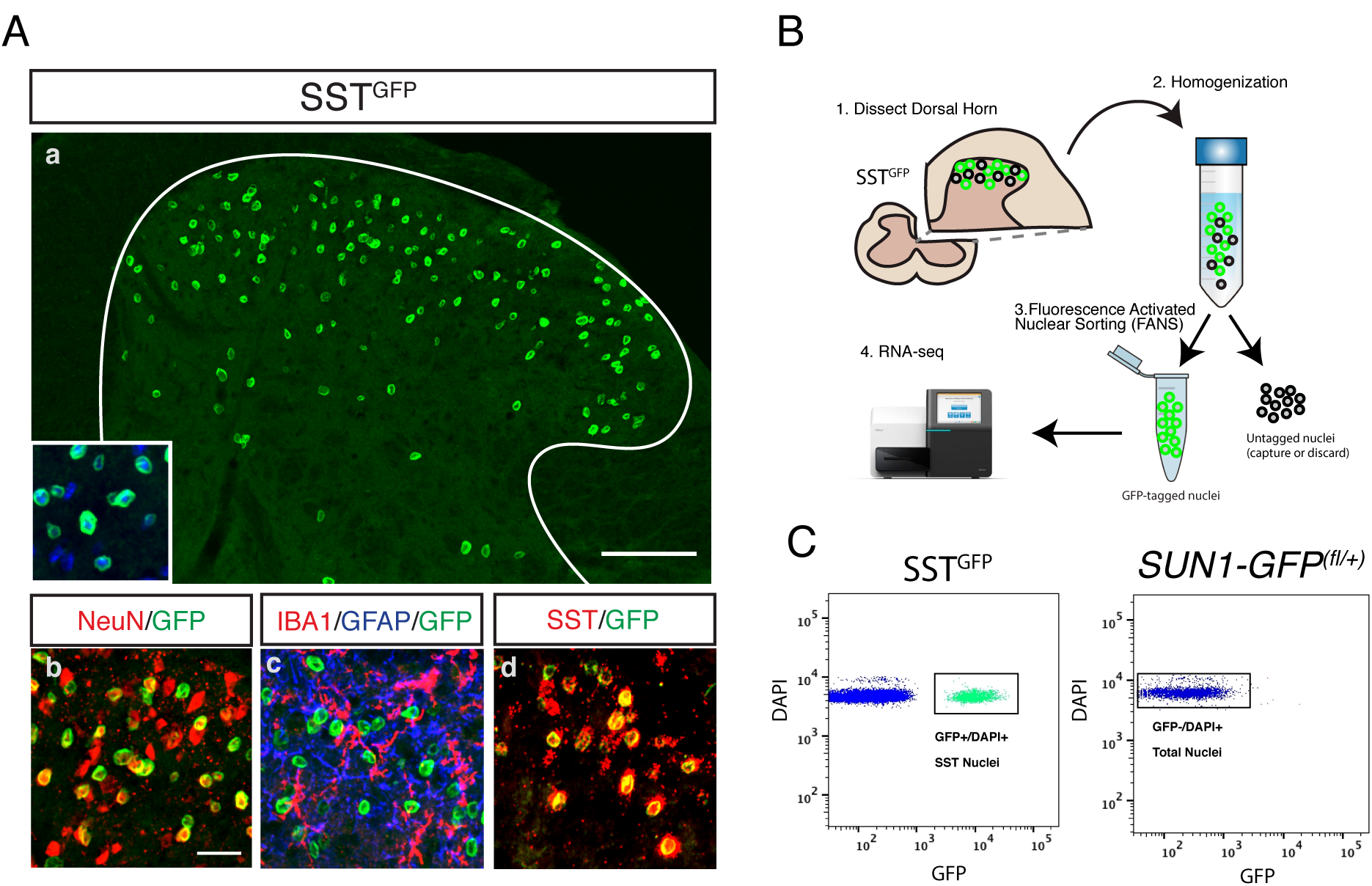
Cell type-specific nuclear tagging and capture of SST neurons in the spinal dorsal horn. (A) SST^GFP^ mice express the Sun1-GFP nuclear tag in SST neurons in the lumbar dorsal horn (a). The GFP-tagged SST neurons co-localize with the neuronal marker NeuN (red, b), but not with the microglial marker IBA1 (red) or the astrocyte marker GFAP (blue) (c). (d) GFP+ (green) nuclei overlap extensively with native *Sst* mRNA (red). (B) Schematic of overall experimental workflow. (1) The dorsal segment from one side of the lumbar spinal cord is dissected. (2) Nuclei are liberated by Dounce homogenization in nuclear lysis buffer. (3) Nuclei are then captured using Fluorescence Assisted Nuclear Sorting. (4) The isolated RNA is processed for RNA-seq. (C) GFP+/DAPI+ nuclear were captured from SST^GFP^ (left). No GFP+ nuclei were present in Sun1-GFP^*(fl/+)*^ control animals; all DAPI+ nuclei were captured.

Combined immunostaining and *in situ* hybridization showed that that GFP+ nuclei co-localized extensively with native *Sst* mRNA expression (81.8% ± 2.4%, n=5), as well as with the neuronal marker NeuN, but not with GFAP or IBA1, markers of astrocytes and microglia, respectively (Figure 1Ab-d). Staining with the nuclear dye DAPI showed nuclear GFP signal localized at the periphery of the DAPI+ nucleus (Figure 1Aa, inset).

Having established that the SST^GFP^ mouse faithfully labels the majority of Sst-expressing neurons in the dorsal horn, we used Fluorescence Activated Nuclear Sorting (FANS) to selectively collect GFP+ nuclei from the lumbar dorsal segment of 8-week old SST^GFP^ mice (Figure 1B) (Krishnaswami et al., 2016). To distinguish nuclei from cellular debris, DAPI was used to stain all intact nuclei. Five thousand (5000) GFP+/DAPI+ nuclei were sorted (n=3) directly into lysis buffer. For comparison, the same number of total dorsal horn nuclei (DAPI+/GFP-) was captured from Cre-negative Sun1-GFP^(fl/+)^ control mice (n=3)(Figure. 1C). Total RNA was isolated from the samples and RNA-seq libraries were generated.

### Transcriptional Profiling of SST neurons

RNA-seq was performed on the libraries generated from spinal nuclei. Sequencing depth per sample was 25.8 million reads (average). We detected 13,342 genes (coding and non-coding) using a normalized count cutoff of 10 (Table 2-1). Pearson correlation hierachical clustering indicated that the transcriptional profiles of the SST^GFP^ population differed markedly from those of the total population (Figure 2A). To identify specific genes enriched in SST neurons, we performed differential gene expression analysis, comparing the SST^GFP^ profiles to those of the total nuclei. We observed 901 protein-coding, differentially expressed genes (DEGs) with a log2-fold change (log2FC) > 1, counts >20, and an adjusted p-value (q-value) < 0.01. SST neurons were greatly depleted of glial markers such as *Aldh1l1, Cx3cr1* and *Mog*, representing astrocytes, microglia and oligodendrocytes, respectively (Figure 2B). Among the top 50 DEGs (Figure. 2C, Table 1), numerous genes with known expression in the dorsal horn and roles in nociception were detected, supporting the validity of both the technique and our results. Examples of known pain and dorsal horn genes include *Nmur2, Nmu, Tlx3, Prrxl1, Tac1, Lmx1b, Prkcg, Cacna2d1* and *Grp* (LaCroix-Fralish et al., 2007). Numerous novel genes without previous description in the dorsal horn also appeared among the top 50 DEGs in SST neurons, several of which we validated and describe below.

**Figure 2.**
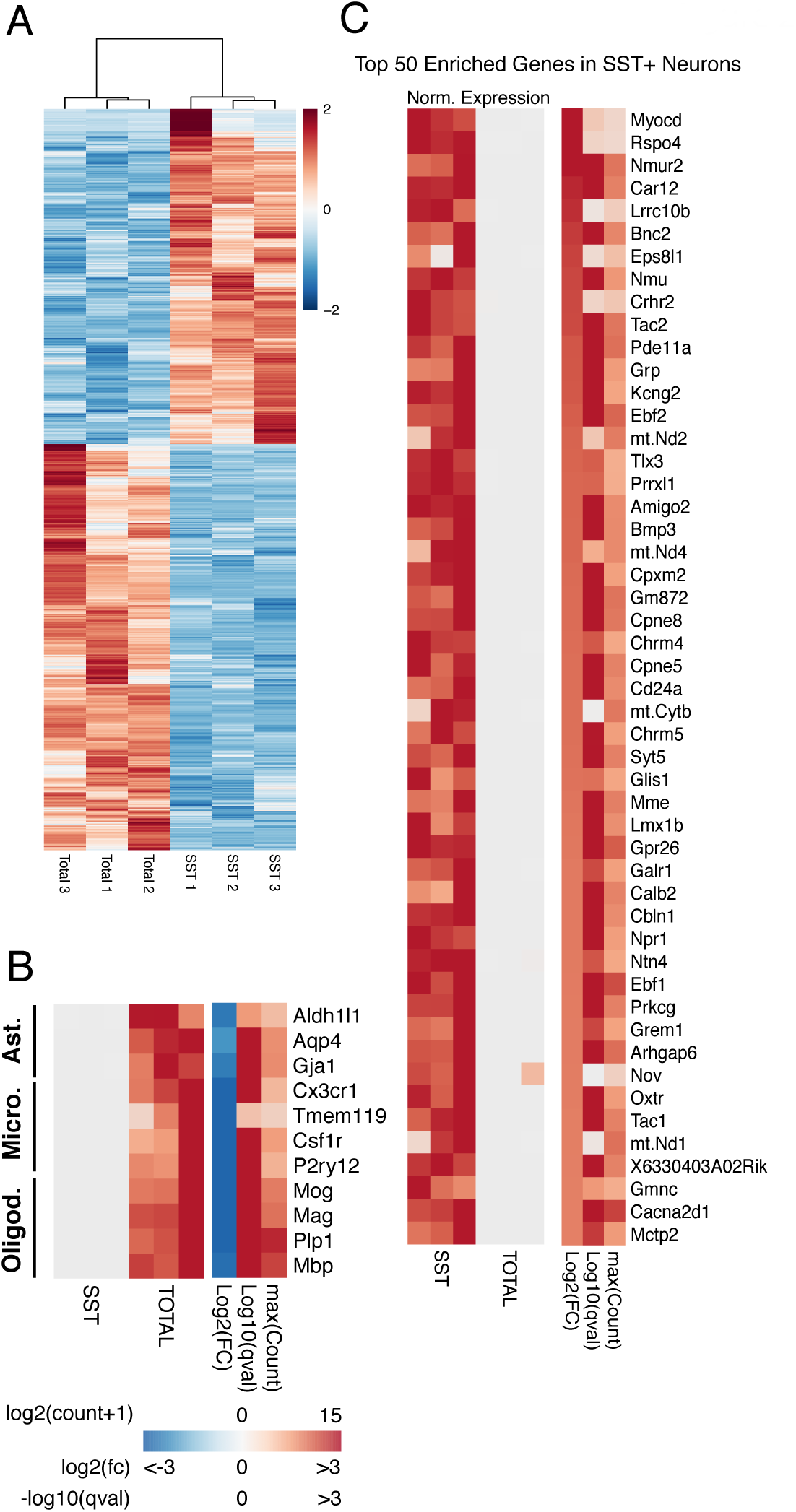
Differential Gene Expression in SST Neurons. (A) Hierarchical clustering of SST neurons and total dorsal horn nuclei reveals widely divergent gene expression patterns. Color scale indicates Z-score of normalized expression values. (B) Non-neuronal markers are depleted from SST neurons. Ast. = Astrocytes, Micro.= Microglia, Oligod. = Oligodendrocytes. In the left heatmap, the expression level in each sample is normalized by the maximum expression level in the row (Norm. Expression). The right heatmap shows the log2-Fold Change (log2FC),-log10 of the q-value (adjusted p-value) and the log2 of the maximum value of counts for each gene plus a psuedocount of 1 (maxCount). (C) Top 50 Differentially Expressed Genes in SST neurons. Genes are ordered by log2FC value from highest to lowest. Criteria for inclusion of genes: log2FC > 1, q-value <0.01 and CV <65% and a minimum of 20 counts for all samples. Scale

**Figure 3.**
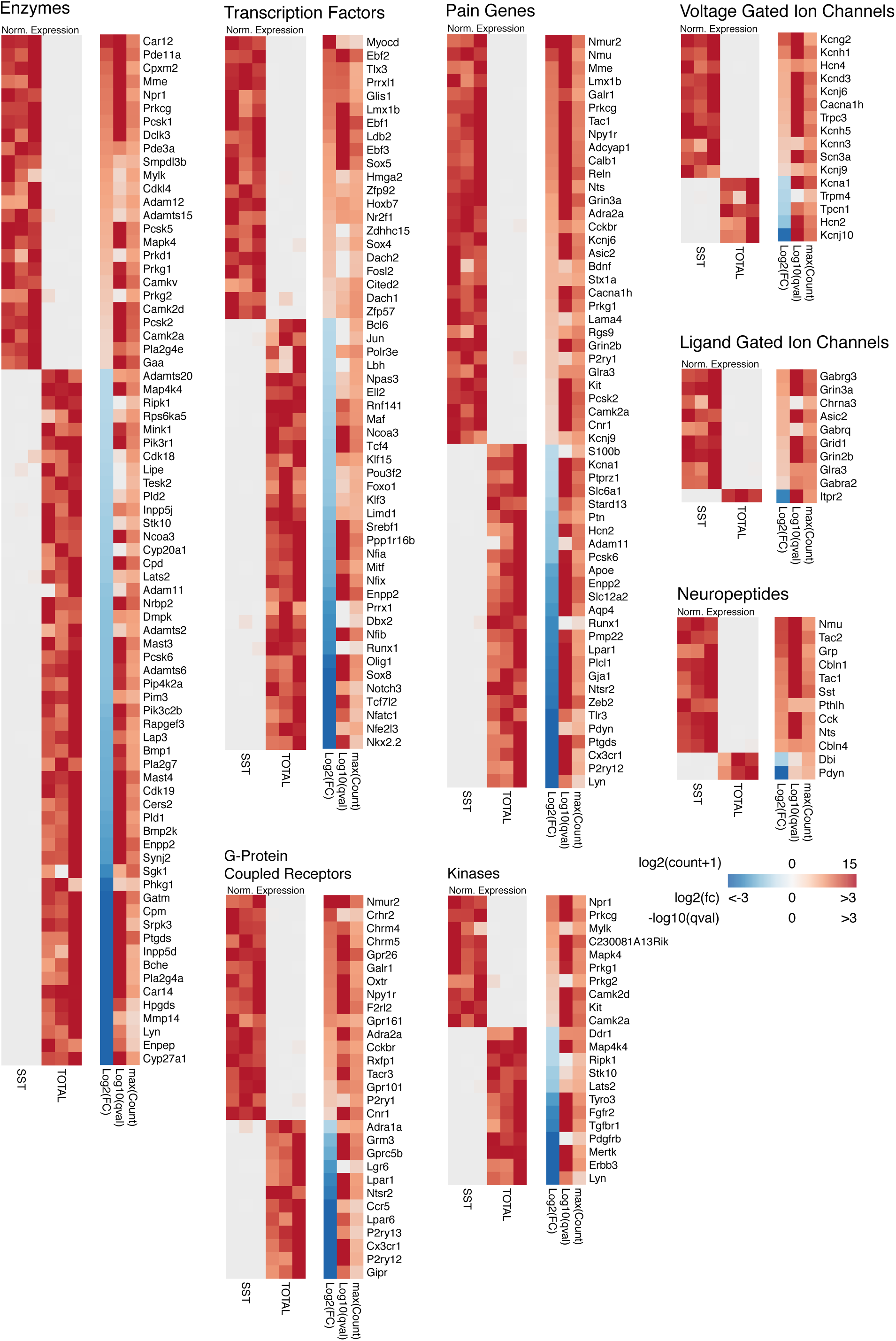
Enrichment of Genes from Selected Functional Classes in SST Neurons. Differential expression of genes in SST neurons compared to total dorsal horn nuclei grouped by functional classes. For each class, in the left heatmap, the expression level in each sample is normalized by the maximum expression level in the row (Norm. Expression). The right heatmap shows the log2-Fold Change (log2FC),-log10 of the q-value (adjusted p-value) and the log2 of the maximum value of counts for each gene plus a psuedocount of 1 (maxCount).

**Table 1.** Top 50 differentially expressed genes in SST neurons compared to total dorsal horn cells. For each gene we report: gene, log2(fold change), Wald Test Statistic, q-value (adjusted p-value) and the log2(Max Count), which is log base-2 of the maximum normalized count value. n=3/group.

| <u>Gene</u> | <u>log2FC</u> | <u>Wald Test Statistic</u> | <u>q-value</u> | <u>log2 (Max Count)</u> |
| --- | --- | --- | --- | --- |
| Myocd | 3.5 | 5.2 | 2.2E-06 | 4.9 |
| Rspo4 | 3.3 | 4.9 | 8.1E-06 | 4.8 |
| Nmur2 | 3.1 | 10.0 | 8.4E-22 | 10.1 |
| Car12 | 2.8 | 17.3 | 1.6E-64 | 9.4 |
| Lrrc10b | 2.8 | 4.5 | 5.6E-05 | 5.4 |
| Bnc2 | 2.6 | 14.3 | 3.0E-44 | 9.2 |
| Eps8l1 | 2.6 | 4.8 | 1.8E-05 | 6.1 |
| Nmu | 2.5 | 12.0 | 4.2E-31 | 8.2 |
| Crhr2 | 2.5 | 4.9 | 8.6E-06 | 5.8 |
| Tac2 | 2.5 | 17.1 | 7.7E-63 | 10.0 |
| Pde11a | 2.4 | 17.1 | 2.2E-63 | 10.4 |
| Grp | 2.3 | 8.7 | 7.8E-17 | 8.0 |
| Kcng2 | 2.3 | 9.1 | 2.6E-18 | 7.5 |
| Ebf2 | 2.3 | 17.7 | 2.2E-67 | 10.9 |
| mt-Nd2 | 2.2 | 5.3 | 1.6E-06 | 9.7 |
| Tlx3 | 2.2 | 7.3 | 4.4E-12 | 7.0 |
| Prrxl1 | 2.2 | 7.2 | 1.1E-11 | 7.1 |
| Amigo2 | 2.2 | 14.0 | 2.5E-42 | 8.9 |
| Bmp3 | 2.2 | 12.6 | 1.5E-34 | 8.9 |
| mt-Nd4 | 2.2 | 5.8 | 9.2E-08 | 8.9 |
| Cpxm2 | 2.1 | 9.2 | 1.9E-18 | 7.7 |
| Gm872 | 2.1 | 14.2 | 1.9E-43 | 10.1 |
| Cpne8 | 2.1 | 16.0 | 2.9E-55 | 10.0 |
| Chrm4 | 2.1 | 7.4 | 2.1E-12 | 7.3 |
| Cpne5 | 2.1 | 12.9 | 5.4E-36 | 9.4 |
| Cd24a | 2.1 | 12.4 | 3.8E-33 | 8.9 |
| mt-Cytb | 2.1 | 4.3 | 1.5E-04 | 10.0 |
| Chrm5 | 2.1 | 8.4 | 1.1E-15 | 7.8 |
| Syt5 | 2.1 | 13.4 | 1.0E-38 | 9.5 |
| Glis1 | 2.0 | 7.0 | 4.4E-11 | 7.4 |
| Mme | 2.0 | 10.6 | 2.1E-24 | 9.6 |
| Lmx1b | 2.0 | 9.5 | 1.1E-19 | 8.7 |
| Gpr26 | 2.0 | 21.8 | 4.0E-102 | 11.4 |
| Galr1 | 2.0 | 7.7 | 4.2E-13 | 7.7 |
| Calb2 | 2.0 | 9.0 | 9.1E-18 | 9.3 |
| Cbln1 | 2.0 | 12.2 | 4.9E-32 | 8.8 |
| Npr1 | 2.0 | 8.9 | 1.6E-17 | 7.9 |
| Ntn4 | 2.0 | 7.5 | 1.6E-12 | 7.3 |
| Ebf1 | 2.0 | 21.7 | 9.5E-102 | 12.2 |
| Prkcg | 1.9 | 13.1 | 4.5E-37 | 9.5 |
| Grem1 | 1.9 | 7.7 | 2.6E-13 | 7.6 |
| Arhgap6 | 1.9 | 14.7 | 1.1E-46 | 10.5 |
| Nov | 1.9 | 3.7 | 1.2E-03 | 5.2 |
| Oxtr | 1.9 | 9.1 | 2.2E-18 | 8.0 |
| Tac1 | 1.9 | 13.2 | 1.7E-37 | 9.4 |
| mt-Nd1 | 1.9 | 4.5 | 6.1E-05 | 10.3 |
| 6330403A02Rik | 1.9 | 13.2 | 1.9E-37 | 9.4 |
| Gmnc | 1.9 | 6.2 | 7.4E-09 | 7.1 |
| Cacna2d1 | 1.9 | 18.7 | 3.2E-75 | 12.8 |
| Mctp2 | 1.9 | 7.9 | 5.9E-14 | 8.0 |

### Functional Classes of Genes Expressed by SST neurons

To further examine enrichment of genes that are pharmacologically tractable, we grouped DEGs into pharmacologically relevant functional classes.

### Enzymes

*Prkcg* encodes for Protein Kinase C gamma isoform (PKCγ), which marks a class of excitatory interneuron involved in mechanical allodynia and contributes to the intracellular signaling cascade that underlies this role (Neumann et al., 2008). Previous histological studies have shown that PKCγ colocalizes with approximately one-third of SST-expressing interneurons in lamina I-II (Gutierrez-Mecinas et al., 2016). Consistently, *Prkcg* is one of the most highly enriched enzymes and is among the top 50 DEGs overall (+1.9 log2FC) in SST neurons.

*Mme* encodes for membrane metallo-endopeptidase, also known as neprilysin, which has been implicated in pain processing via its role in the breakdown of enkephalins and other peptides (Krämer et al., 2009) *In situ* hybridization studies showed strong and specific expression in the superficial (laminae I-II) dorsal horn of the rat, but more precise characterization of its expression is lacking (Facchinetti et al., 2003). Our results here indicate that *Mme* is enriched in SST neurons (+2.0 log2FC). Given the known function of MME in regulating endokephalin levels and the function of SST neurons in mechanical pain, MME may be used by SST neurons to enhance noxious mechanical transmission by downregulating endogenous opioids.

Within the enzyme class, we also discovered several novel targets with no previously described role in pain or expression in the dorsal horn.

*Car12* encodes for Carbonic Anhydrase 12, which is a membrane spanning carbonic anhydrase whose catalytic domain faces the extracellular space (Türeci et al., 1998). A recent large scale single-cell RNA-seq study in the visual cortex identified *Car12* as a unique marker of Layer 6 cortical neuron subtype (Tasic et al., 2016), but apart from this, virtually nothing is known about the function of *Car12* in the nervous system. However, other extracellular carbonic anhydrases (ECAs) such as *Car4* and *Car14* have been well-studied in the brain, where they have been shown to modulate neuronal activity by regulating extracellular pH (Makani and Chesler, 2007; Svichar et al., 2009; Makani et al., 2012). In the hippocampus, several studies have shown that ECAs regulate excitatory synaptic transmission via pH-mediated modulation of NMDA receptor activation(Chen and Chesler, 2015). Given that the majority of the SST neurons in the dorsal horn are excitatory, the distinct enrichment of *Car12* in these neurons may indicate that it regulates NMDA-mediated excitatory synaptic transmission in a manner analogous to that observed in the hippocampus.

To validate the expression of *Car12*, we performed both *in situ* hybridization and immunostaining. *In situ* hybridization showed strong expression of *Car12* transcripts exclusively in the superficial dorsal horn (Figure 4Aa) that co-localized extensively with *Sst* transcript as well as GFP+ nuclei (Figure 4Ab-e). Nearly half of all cells expressing *Sst* transcript expressed *Car12* (46.9% ± 0.7%, n=5), and the majority of Car12-expressing neurons expressed *Sst (69.2%* ±0.1%, n=5). 42.7% ± 2.1% of GFP+ neurons co-expressed both *Car12* and *Sst* mRNA. Because *Car12* is an ECA, it is expected to localize to the plasma membrane of the cells in which it is expressed. To better visualize the soma of SST neurons, we employed an SST-tdTomato mouse, which expresses cell-filling tdTomato (Duan et al., 2014). Immunostaining with an antibody against mouse CAR12 showed CAR12-immunoreactivity (CAR12-IR) localized on the membrane of SST-Tomato neurons in superficial laminae, consistent with the findings for Car12-mRNA (Figure 3Ae-g).

**Figure 4.**
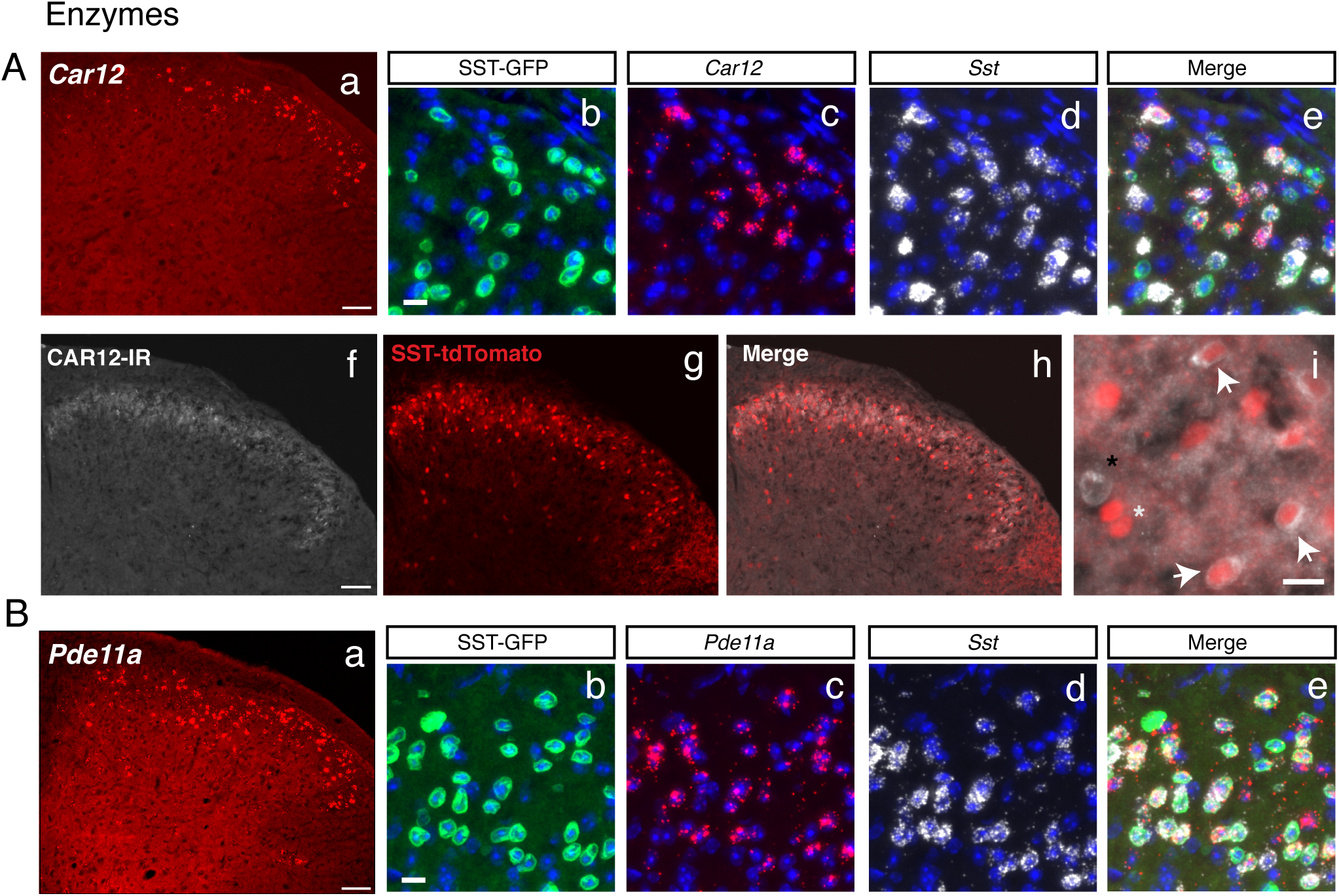
Expression of the enzymes *Car12* and *Pdella* in SST neurons in the dorsal horn. (A) *(a)Car12* (red) *in situ* hybridization in the dorsal horn at low magnification (20x). Scale = 50 μm. High magnification (40x) image of GFP (b), *Car12* (c), *Sst* (d) and merge (e). DAPI (blue) in images. Scale = 10 μm. (f) Immunohistochemistry of CAR12-IR, (g) SST-tdTomato and (h) merge at magnification (20x). Scale = 100 μm. (i) High magnification (40x) image of SST-tdTomato and CAR12-IR. Scale = 10 μm. White arrows indicate cells with tdTomato and CAR12-IR colocalization. White asterisk indicates tdTomato+ cells that lack CAR12-IR and black asterisk indicates a CAR12-IR+ without tdTomato. (B) (a) *Pdella* (red) *in situ* hybridization in the dorsal horn at low magnification (20x). Scale = 50 μm. High magnification (40x) image of GFP (b), *Pdella* (c), *Sst* (d) and merge (e). DAPI (blue) in images. Scale = 10 μm.

Phosphodiesterase 11A (PDE11A) is a dual specificity phosphodiesterase that catalyzes the breakdown of the cyclic nucleotides cAMP and cGMP. PDE11A exhibits restricted expression in the brain, predominating in the ventral hippocampus(Kelly et al., 2010). Knockout studies of PDE11A have implicated this gene in regulating social behaviors and anxiety(Hegde et al., 2016a; 2016b). In the dorsal horn, cyclic nucleotide signaling contributes to pain-related neuronal plasticity, and other members of the phosphodiesterase family have been directly implicated in the pathogenesis of inflammatory and neuropathic pain(Kallenborn-Gerhardt et al., 2014). Previous studies have detected *Pde11a* transcript and PDE11A protein in the bulk analyses of the spinal cord, but its cellular expression pattern is unknown(Kelly, 2015). Accordingly, we selected *Pde11a* for further examination. *In situ* hybridization of *Pde11a* transcript revealed a strong and specific signal that predominated in the superficial laminae (Figure. 4Aa). Half of all Sst-expressing neurons co-expressed *Pde11a* (50.8% ± 1.3, n=3), and conversely, nearly two-thirds of Pde11a-expressing neurons co-expressed *Sst* mRNA (64.2% ± 1.9%, n=3). More than half of all GFP+ nuclei in the dorsal horn co-localized with both *Pde11a* and *Sst* mRNA (55.8% ± 3.0%, n=3). As with *Car12*, the restricted co-expression of *Pdella* and *Sst* in only the most superficial laminae suggests that this gene marks a functional subtype of Sst-expressing neurons.

### Transcription Factors

Transcription factors are responsible for the specification of neuronal identity(Ma, 2006). Much work has been conducted on the role of specific transcription factors in the development of dorsal horn neurons involved in pain and itch (Ding, 2004; Xu et al., 2013). Nearly all of the well-studied transcription factors that specify excitatory interneuron development were enriched in SST neurons, including *Tlx3, Lmx1b, Prrxl1, Sox5*, and *Ebf1-3* (Chen et al., 2001; Ding, 2004; Li et al., 2006), in support of the biological accuracy of our RNA-seq results. Enriched transcription factors without known functions in dorsal horn interneurons included *Myocd* (+3.5 log2FC), *Glis1* (+2.0 log2FC) and *Ldb2* (+1.7 log2FC) (Li et al., 2006). In addition, a zinc-finger-containing protein, Basonuclin 2 *(Bnc2)* was also among the most highly enriched genes overall. While not a classical transcription factor, by virtue of its zinc finger domain and nuclear localization signal, *Bnc2* regulates transcriptional events(Vanhoutteghem et al., 2016). A recent report that used a LacZ reporter mouse demonstrated that in the central nervous system, *Bnc2* is only expressed in a small subset of dorsal horn cells (Vanhoutteghem et al., 2016), which is consistent with our transcriptomic results. This highly restricted expression of *Bnc2* suggests that it may perform a key function in some SST neurons. Manipulative studies of these undescribed transcription factors would likely reveal important roles in the determination of SST interneuron fates and pain processing.

### G-Protein Coupled Receptors

G-protein coupled receptors (GPCRs) are one of the most important classes of drug targets, making up nearly half of all human drug targets(Bull and Doig, 2015). GPCRs play diverse roles in neurons, functioning prominently as neurotransmitter and neuromodulator receptors(Huang and Thathiah, 2015). Numerous GPCRs that have been implicated in pain processing in the spinal cord were significantly enriched in the SST population, including *Nmur2, Galr1, Npy1r*, and *Cckbr.* In support of our transcriptomic results, previous reports have shown that *Galr1* and *Npy1r* are preferentially expressed in excitatory dorsal horn neurons under the control of *Tlx3(Guo* et al., 2012). Interestingly, two members of the muscarinic acetylcholine receptor family, *Chrm4* (+2.1 log2FC) and *Chrm5* (+2.1 log2FC), were also significantly enriched. As cholinergic signaling is well-known to contribute to pain modulation in the spinal cord, these two receptors may play particular roles in SST neurons gating mechanical pain (Naser and Kuner, 2017).

*Nmur2* is one of two G-protein coupled receptors that bind to the pleotropic neuropeptide neuromedin (NMU). Knockout studies of NMUR2 demonstrated that this receptor contributes to nociception(Torres et al., 2007). Injection of intrathecal NMU enhances synaptic transmission as measured by electrophysiological studies, and produces mechanical and thermal hyperalgesia in behavioral assays (Yu et al., 2003a; Nakahara et al., 2004). Apart from its presence in the dorsal horn, further characterization of the neurons that express *Nmur2* has not been performed. As the third most enriched gene in our top 50 DEGs (Figure 2C), *Nmur2* is one of the more distinctive genes observed, suggesting that it is uniquely expressed by SST neurons. Using *in situ hybridization*, we found *Nmur2* to be expressed nearly exclusively in SST neurons (Figure 4B). As with other genes we examined, only a portion of the *Sst*-expressing neurons co-express *Nmur2*, indicating that these neurons are a unique subpopulation within the broader SST neurons.

We also identified several novel GPCRs whose presence in the dorsal horn has not been described.

*F2rl2* encodes for the protein PAR3, which is a G-protein coupled receptor in the family of Protease Activated Receptors (PARs), of which there are four (PAR1-4) (Saito and Bunnett, 2005). As their names suggest, PARs are activated when proteases such as thrombin or trypsin cleave a portion of the receptor, creating a tethered ligand that then acts as an intramolecular agonist for the receptor. PAR1, 2 and 4 have all been implicated in pain, with PAR2 being the most well-studied(Vellani et al., 2010; Tillu et al., 2015; Mrozkova et al., 2016). PAR3’s function, however, remains elusive, with neither the ligand nor mode of action of this receptor known. In light of the role of the other PARs in pain and itch, the enrichment of PAR3 in SST neurons is suggestive. Thus, we confirmed the expression of PAR3 by *in situ* hybridization. *F2rl2* mRNA strongly expresses in the superficial dorsal horn and co-localizes with the majority of Sst-expressing neurons in the dorsal horn (59.6% +/-2.0%, n=5). More than half of all *F2rl2-*expressing neurons also express *Sst* mRNA (57.1% +/-0.8%, n=5). Future functional studies using knockout mice or pharmacological tools will be needed to determine the role of this gene in spinal nociception.

*Gpr26* is an orphan G-protein coupled receptor that is coupled to G_s_ and for which an endogenous ligand is not known(Khan and He, 2017). Studies with *Gpr26* knockout mice have implicated this receptor in anxiety, depression-like behaviors and obesity, but no known role in pain is known (Zhang et al., 2011). As one of the most enriched GPCRs (+1.9 log2FC) and one of top 50 DEGs (Figure. 2C), we examined the expression of *Gpr26* using *in situ* hybridization. We found this receptor broadly expressed in the dorsal horn, and co-localized with a subset of SST neurons (Figure. 4C). To our knowledge, this is the first report of *Gpr26* in the spinal cord.

### Kinases

Kinases play key roles in intracellular signaling cascades and are frequent targets for pharmacological modulation. Several kinases were enriched in the SST population. *Prkcg*, which has a known role in pain, was the most highly enriched and most distinctive. Other moderately enriched kinases include members of the Calcium-calmodulin Kinase family *(Camk2a* and *Camk2b)*, cyclic-GMP-dependent kinase family *(Prkg1* and *Prkg2)* and Mitogen Activated Protein Kinase family *(Mapk4).*

### Ion Channels

Our RNA-seq results uncovered several previously described ion channels that express in the dorsal horn. *Trpc3* is a TRP channel family member that has been shown to be enriched in the dorsal horn, but no further investigation has characterized its localization or function in spinal neurons(Li et al., 2006). *Hcn4*, which is a hyperpolarization-activated cyclic nucleotide-gated channel, has been shown to colocalize with PKCγ interneurons in lamina IIi, some of which are *Sst*-expressing(Hughes et al., 2013). *Cacna2d1*, which is the target of the clinically effective gabapentinoids, was also highly enriched in SST neurons and with nearly the highest normalized counts of all genes analyzed(Table 1). Many studies of *Cacna2ad1* in pain have examined presynaptic mechanisms in sensory afferents(Taylor and Garrido, 2008; Boroujerdi et al., 2011), but the high expression of this channel subunit in SST neurons suggests a role in these spinal neurons as well. Several ligand-gated ion channels were also enriched, including *Grin3a*, which is an NMDA-receptor subunit known to express in excitatory dorsal horn neurons (Guo et al., 2012) and the acid-sensing *Asic2* channel, which has also been described in dorsal horn neurons(Wu et al., 2004).

Among previously uncharacterized ion channels, the *Kcng2* channel was the most highly enriched in SST neurons. *Kcng2* encodes a gamma subunit of the Kv6 family of voltage-gated potassium channels (Kv6.2). Little is known about the functional role of *Kcng2.* It is highly expressed in myocardial tissue, and so a role in cardiac function has been proposed (Zhu et al., 1999), but no study of *Kcng2* has been conducted in the nervous system. *Kcng2* is enriched in SST neurons (+2.3 log2FC) compared to all cells in the dorsal horn, suggesting a selective role for this channel in affecting pain behaviors by regulating the firing properties of Sst-expressing neurons. In support of this possibility, a recent human Genome-Wide Association Study (GWAS) found human KCNG2 to be significantly associated with opioid dependence.

### Neuropeptides

Neuropeptides play key roles as neuromodulators and often define different populations of neurons, such as is the case with *Sst*. We found several neuropeptides to be enriched in SST neurons. Many of these neuropeptides have been identified and investigated in the dorsal horn, including *Tac2, Grp* and *Sst.* As expected, the defining neuropeptide, *Sst* is enriched (+1.8 log2FC). Also, in further support of our methodology, the neuropeptide *Pdyn*, which is exclusively expressed by inhibitory dorsal horn neurons (Kardon et al., 2014), was strongly depleted in SST neurons (-3.8 log2FC).

The neuropeptide Neuromedin U (NMU) is encoded by the *Nmu* gene and exerts wide-ranging effects both in the periphery and CNS. NMU is abundant in the spinal cord and knockout of *Nmu* in the mouse produces large deficits in the hot plate and formalin tests(Nakahara et al., 2004). Exogenous NMU enhances nociceptive behaviors, including mechanical sensibility(Yu et al., 2003b). The cellular expression profile of *Nmu* in the spinal cord is unknown. Our RNA-seq data indicate that *Nmu* is enriched (+2.5 log2FC) in SST neurons, suggesting that some SST neurons use this neuropeptide in nociceptive signaling. As we demonstrated, the receptor for NMU, *Nmur2*, was also highly enriched in a subset of Sst-expressing neurons in the superficial dorsal horn (Figure. 4B). Whether NMU and NMUR2 are co-expressed in the same neurons will need further study, but our findings suggest a possible autocrine mechanism, or intercellular signaling between different subsets of SST neurons.

*Tac2* encodes the neuropeptide Neurokinin B(Gutierrez-Mecinas et al., 2016). *Tac2* was highly enriched in the SST population. This neuropeptide has been identified in dorsal horn neurons and found to overlap in part with PKCγ-expressing neurons, some of which are also *Sst-*expressing. However, toxin-mediated ablation of the *Tac2*-lineage neurons in the spinal cord did not affect mechanical pain behaviors(Duan et al., 2014), in contrast to the dramatic effect of ablation of the SST neurons. This indicates that the subset of SST neurons that expresses *Tac2* is dispensable for mechanical pain behaviors. What role these *Tac2/Sst* co-expressing neurons plays in pain remains to be determined.

*Grp* encodes the Gastrin Related Peptide (GRP), a neuropeptide that has been well-studied for its role in itch transmission in the dorsal horn(Mishra and Hoon, 2013; Hoon, 2015). Recent work has demonstrated that GRP is expressed by a subset of dorsal horn interneurons and is used for pruriceptive transmission. Moreover, it is known that these GRP-expressing interneurons co-express the Natriuretic Peptide Receptor A *(Npr1)* and release GRP when NPR1 is activated by its cognate ligand Natriuretic Polypeptide B (Nppb) (Mishra and Hoon, 2013). Our RNA-seq data indicate that both *Grp* (+2.5 log2FC) and *Npr1* (+2.0 log2FC) are enriched in SST neurons, suggesting that the pruriceptive *Grp+/Npr1+* population is subsumed in the SST population. Consistently, a recent study demonstrated that optogenetic activation of SST neurons in the dorsal horn elicited strong light-induced itch behaviors(Christensen et al., 2016). Another study used a chemogenetic approach to manipulate Grp-expressing spinal neurons and found that this population mediates both pain and itch in an intensity-dependent manner(Sun et al., 2017). Taken together, our transcriptomic findings along with these manipulative studies suggests that an interneuron population expressing *Gpr/Sst/Npr1* may exist that can mediate both mechanical pain and itch.

### Other Genes

Other genes with high enrichment and relatively high counts appeared that do not fall into the functional classes described above (Table 1). *Bmp3* encodes Bone Morphogenetic Protein 3, which is a secreted ligand of the TGF-β family. *Bmp2, Bmp4* and *Bmp7* play critical roles in the specification of dorsal spinal neurons, but no role for *Bmp3* has been described, although the expression of BMP3 in the adult dorsal horn by immunohistochemistry has recently been described (Yamashita et al., 2016). As a secreted ligand, BMP3 may function in intercellular signaling in the dorsal horn.

*Cpne5* and *Cpne8* encode for Copine 5 and Copine 8, respectively. Copines are a family of calcium-dependent phospholipid-binding proteins. Copine 6 has been shown to regulate excitatory synapses in the hippocampus by translating calcium signals into changes in dendritic spine structure through Rac1 recruitment (Reinhard et al., 2016). Copine 5 and Copine 8 may function analogously in regulating the excitatory synapses of SST neurons in the dorsal horn and warrant further study.

### Pathway Analysis

Genes that participate in functional pathways are often coordinately expressed. To determine whether known signaling pathways are enriched in SST neurons, we applied Ingenuity Pathway Analysis. Using the same selection criteria as for single genes (q < 0.01, log2FC > 1), we identified several enriched pathways and ranked them by their statistical significance (Figure 5). Top among the pathways was “Neuropathic Pain in Dorsal Horn Neurons”, which is consistent with the known role of SST neurons in neuropathic pain models. The next identified pathway was cAMP-mediated signaling followed by “nNOS signaling in neurons” and “G-Protein Coupled Receptor signaling”. To complement the Ingenuity analysis, we also applied Gene Set Enrichment Analysis (GSEA), which is used to identify coordinate changes in the expression of groups of functionally related genes (Mootha et al., 2003). From this analysis, the top gene set was Voltage Gated Calcium Channel Activity, followed by Neurotransmitter Receptor Activity and Calcium Channel Activity. These gene set enrichments suggest that calcium signaling is particularly important in the SST neurons.

**Figure 5.**
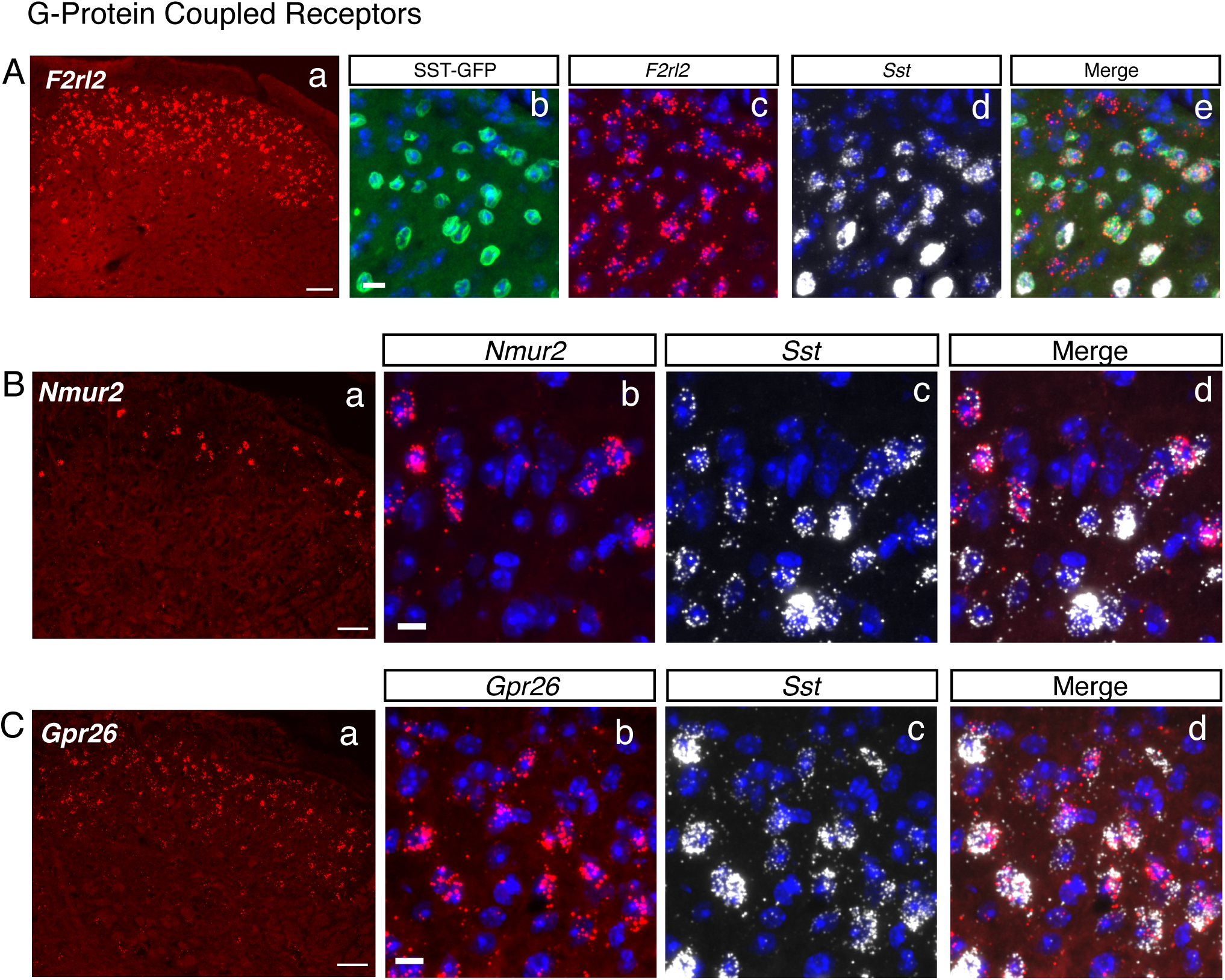
Expression of the G-protein Coupled Receptors *F2rl2, Nmur2* and *Gpr26* in the dorsal horn. (A) (a) *F2rl2* (red) *in situ* hybridization in the dorsal horn at low magnification (20x). Scale = 50 μm. High magnification (40x) image of GFP (b), *F2rl2* (c), *Sst* (d) and merge (e). DAPI (blue) in images. Scale = 10 μm. (B) (a) *Nmur2* (red) *in situ* hybridization in the dorsal horn at low magnification (20x). Scale = 50 μm. High magnification (40x) image of *Nmur2* (b), *Sst* (c) and merge (d). DAPI (blue) in images. Scale =10 μm. (C) (a) *Gpr26* (red) *in situ* hybridization in the dorsal horn at low magnification (20x). Scale = 50 μm. High magnification (40x) image of *Gpr26* (b), *Sst* (c) and merge (d). DAPI (blue) in images. Scale = 10 μm.

**Figure 6.**
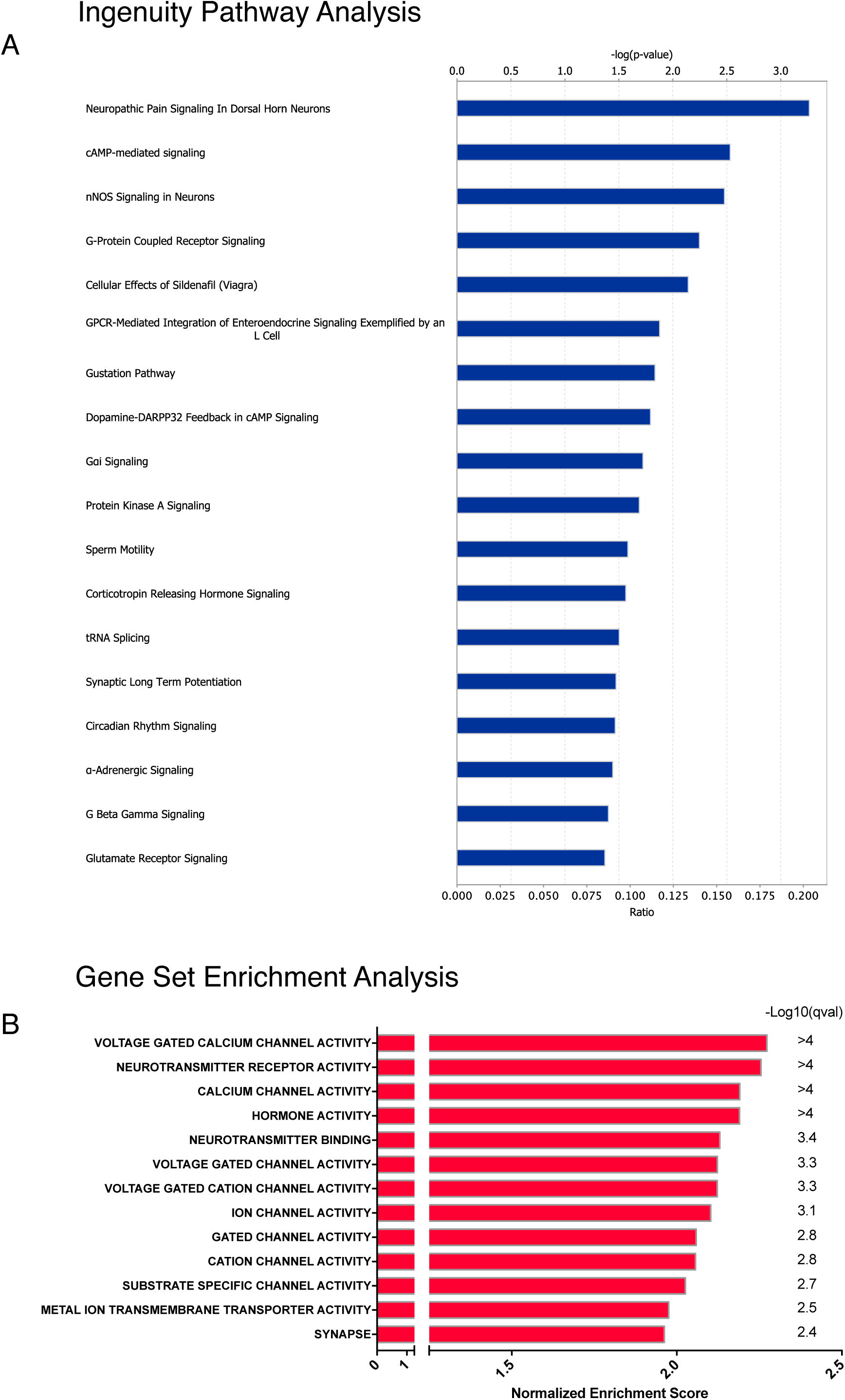
Pathway analysis and Gene Set Enrichment Analysis of SST neurons. (A) Ingenuity Pathway Analysis reveals multiple enriched pathways in SST neurons. Top axis = – log(p-value). Bottom axis = ratio of enrichment in canonical pathway. (B) Gene Set Enrichment Analysis of SST neurons.

## Discussion

In this study we present a comprehensive transcriptomic profile of SST neurons in the spinal dorsal horn of the mouse and identify several novel differentially expressed genes that could be attractive therapeutic targets and population markers.

We optimized the INTACT method for spinal tissue in order to profile the transcriptome of SST neurons in the dorsal horn. To our knowledge, this is the first demonstration of cell type-specific transcriptomic profiling of a neuron population from this region, and the first to use nuclei in lieu of cells. Our results show that neuronal nuclei are suitable surrogates for whole cells for RNA-seq profiling of neuronal populations of the dorsal horn. Indeed, given the difficulty of obtaining viable whole neurons (YQ, unpublished data), nuclei may be the only feasible option for cell type-specific genomic studies in the dorsal horn of adult animals. Thus, our study represents an important technical advance that will enable future interrogation of specific dorsal horn cell populations. Because of the many benefits of nuclei over whole cells, INTACT would be ideal for examining gene expression changes in distinct cell populations in the setting of pathological pain conditions (e.g inflammatory or neuropathic models. Previous gene expression profiling studies have looked at bulk RNA from spinal cord tissue in such pathological pain models, and the resulting signatures are largely dominated by immune and glial signal(Costigan et al., 2009a). The spinal cord-adapted INTACT method we present here will allow researchers to better understand the transcriptional plasticity of specific cell populations, especially smaller populations whose signal is obscured by the averaging effects of bulk RNA-seq. Indeed, the novel genes we identified in this study have likely gone unnoticed in previous genome-wide expression studies(Li et al., 2006) because they comprise only a small fraction of transcripts in bulk RNA from the spinal cord. By isolating a specific population from all surrounding cells, the signal of such population-enriched genes is unmasked. Moreover, because nuclei carry both DNA and RNA, other genomic applications such as ATAC-Seq, ChIP-Seq or bisulfite sequencing could be profitably applied in tandem with RNA-seq to elucidate pain-related epigenetic plasticity in a cell type-specific manner, which is increasingly appreciated to play key roles in pathological pain (Denk and McMahon, 2012; Denk et al., 2016)

Transcriptomic profiling of SST neurons with INTACT allowed us to identify numerous genes that are enriched in these neurons in the dorsal horn. Some of these enriched genes, such as *Tlx3, Lmx1b, Pkcg, Tac2, Grp, Calb2, Ebf1-3*, and *Nmur2* were already known to express preferentially in excitatory neurons of the dorsal horn, of which the SST population comprises a large proportion (Gutierrez-Mecinas et al., 2016). Our results overlap to a high degree with the catalog of dorsally enriched genes identified through microarray screening by Li *et al.* (Li et al., 2006). Identification of these known targets lends support for the overall biological validity of our findings.

The major aim of this work was to discover novel genes that are enriched in SST neurons. Indeed, we report here several genes whose expression in the dorsal horn was previously unknown and whose annotated functions make them attractive for pharmacological intervention. Using *in situ* hybridization, we characterized the expression of four novel genes that were highly enriched in SST neurons: *Pde11a, Car12, F2rl2* and *Gpr26.* Our results show that each of these genes is expressed in a subset of *Sst*-expressing neurons in the superficial dorsal horn, with *Pde11a* and *Car12* showing particularly restricted expression in laminae I-II. Further studies using specific genetic and pharmacological tools will be needed to determine whether and how each gene contributes to pain processing in *Sst*-expressing neurons.

Our findings also highlight the transcriptional heterogeneity of the SST population and suggest possible subpopulations. Among the most highly enriched genes are markers for multiple neurochemically distinct excitatory neuron populations. In a quantitative study of lamina I-III excitatory interneurons, Gutierrez-Mecinas *et al.* demonstrated that the SST population subsumes three largely non-overlapping excitatory populations defined by *Grp, Tac2* and *Nts*, leading the authors to conclude that SST broadly marks most excitatory interneurons of the superficial dorsal horn(Gutierrez-Mecinas et al., 2016). Our RNA-seq findings are entirely consistent with this study from Gutierrez-Mecinas and support the conclusion that the SST population is comprised of multiple excitatory subpopulations. The restricted expression of *Car12* and *Pde11a* suggests that these genes define subpopulations of excitatory interneurons that may be functionally and neurochemically distinct from other known subtypes. *Nmur2* also appears in a sparse and restricted population that may be distinct as well. Cre driver lines under the control of these genes would be useful to explore the functional role of these populations. *F2rl2* and *Gpr26*, on the other hand, have broader expression, and likely mark most or all excitatory interneurons lamina I-III. Accordingly, these GPCRs would be better suited as pharmacological targets to modulate excitatory synaptic transmission rather than as cell type-markers. The endogenous ligands for and functions of these GPCRs in pain processing remain to be determined.

In addition to looking at single genes we also applied pathway analysis to uncover pathways and functional groups of genes enriched in SST neurons. Using Ingenuity Pathway Analysis, we revealed that SST neurons exhibit enrichment in several druggable pathways including cAMP-, nNOS-and GPCR-mediated pathways. Gene Set Enrichment Analysis revealed an enrichment in calcium channel-related gene sets, pointing toward calcium signaling as an important contributor to the functional properties of SST neurons. Future work that manipulates these pathways as a whole will be needed to understand their contributions to the functional properties of SST neurons.

Our results also highlight some of the limitations of cell type-specific profiling using INTACT. Because INTACT and related cell type-specific methods rely on a single Cre driver line to tag a cell population, the specificity of the readout depends on the specificity of the Cre driver line. As we and others have demonstrated, the SST population is internally heterogeneous, (Duan et al., 2014; Gutierrez-Mecinas et al., 2016). Intersectional methods that use two recombinases (i.e. Cre and Flp) driven by separate genes could be used with INTACT to more precisely access the transcriptomic signatures of discrete dorsal horn neuronal subtypes(Dymecki and Kim, 2007; Duan et al., 2014), but a new intersectional INTACT mouse or viral vector would need to be generated. Additionally, as is true of all transcriptomic methods, mRNA levels do not correlate entirely with protein levels(Maier et al., 2009). These limitations aside, INTACT is a powerful and enabling technology for cell type-specific genomics in the spinal cord.

This study also demonstrates the general utility of nuclei as surrogates for cells in the study of spinal cord transcriptomics. Single-cell genomics is revolutionizing our understanding of cell types in the nervous system(Macosko et al., 2015; Zeisel et al., 2015; La Manno et al., 2016; Habib et al., 2017) and large international efforts are now underway to comprehensively catalog the cells of the mouse and human brain(Ecker et al., 2017). The spinal cord also stands to benefit greatly from these efforts so that a comprehensive census of the neuronal and non-neuronal cell types in the spinal cord can be determined. Based on our experience from this present study and pilot studies using single nuclei from the dorsal horn (AC, YQ, unpublished data), we submit that single nuclei, as opposed to single cells, will provide the most comprehensive and unbiased representation of cell types in the spinal cord, given the difficulties of dissociating viable neurons from adult spinal cord. Indeed, single-nucleus RNA-seq (snRNA-seq) has emerged as an effective strategy in the profiling of single cells in the brain and spinal cord(Habib et al., 2016; 2017) already. Going forward, we propose that single-nucleus RNA-seq and population-level INTACT be used in a complementary manner. Massively parallel snRNA-seq with high numbers of nuclei would be the most effective way to identify and classify cell types in the spinal cord, but would most likely lack the depth to reveal disease or treatment effects on low-to medium-abundance transcripts(Habib et al., 2017). Population-level INTACT and deep sequencing, as we present here, would have the ability to detect transcriptional perturbations from manipulations (e.g. injury, inflammation, drugs) that would be missed by single nuclei.

Overall, our work provides a framework for combining knowledge of functional pain circuitry with transcriptomics to uncover novel genes and to refine our understanding of cell types in the spinal cord.

## Acknowledgements

We thank Ian Cumming at the Duke Human Vaccine Institute Flow Cytometry Core Facility and David Corcoran at the Duke Genomic Analysis and Bioinformatics shared resource for their assistance. We thank Anne West for help and suggestions regarding INTACT. This work was supported by NIH grants 2T32GM008600 and Congressionally Directed Medical Research Programs and the Department of Defense awards W81XWH-12-2-0129 and W81XWH-15-2-0046(TV) and by NIH R01 grants NS87988 and DE17794 to R.R.J. We thank members of the Ji and Van de Ven labs for comments on the manuscript.

---

**Extended Data Table 2-1.** SST and Total Expression Data. This table contains the output of differential expression analysis using DEseq2 with exon-aligning reads. GeneID = Ensembl Gene identification. GeneName = Gene Symbol Name. IsCoding = indicates whether the gene is protein coding. Log2FoldChange = The log_2_ of the fold change between SST and Total. Stat = Wald test statistic value. p-value = unadjusted p-value. padj = p-value adjusted for multiple comparisons, which we refer to in the text as q-value. Columns SST1-3 and TOT1-3 show the normalized counts for a given gene in each biological replicate from the SST neuron set (SST) and Total nuclei set (TOT1-3).

